# Stepping-Stone Population Structure and Widespread Clonality of the Mesophotic Octocoral *Swiftia exserta* in the Warm Temperate Northwest Atlantic

**DOI:** 10.64898/2026.09.01.748044

**Authors:** Nicole C. Pittoors, Luisa Lopera, Samuel A. Vohsen, Matthew P. Galaska, Destiny West, Andrea M. Quattrini, Annalisa Bracco, Santiago Herrera

## Abstract

Isolated mesophotic banks form spatially discrete networks of patchy ecosystems along continental shelves. Their long-term persistence depends on connectivity among populations of the dominant coral species that structure these communities. Yet the scale of larval-mediated gene flow across these networks remains poorly characterized. Similarly, the extent to which clonal propagation shapes local population dynamics is unresolved, leaving the relative contributions of sexual and asexual reproduction to population maintenance an open question. Here, population genomics and high-resolution (1km) biophysical larval dispersal modeling are integrated to resolve the genetic connectivity and clonal dynamics of *Swiftia exserta*, a key habitat-forming octocoral, across 13 mesophotic banks spanning the U.S. Gulf and eastern coast within the Warm Temperate Northwest Atlantic (WTNWA) coastal and shelf biogeographic province. Genome-wide SNP markers derived from Restriction-site Associated Sequencing (RADseq) were genotyped across 288 individuals. About 40% of individuals belonged to clonal genets confined to single banks within meters of one another, indicating that asexual propagation sustains local density but plays no detectable role in inter-bank connectivity. Population genetic analyses pointed to isolation by distance as the primary structuring force, with the majority of molecular variance residing within rather than among populations. The west-most population, East Flower Garden Banks, was the most differentiated population in the WTNWA Northern Gulf ecoregion, reflecting its peripheral position within the bank network. The Edisto population from the WTNWA Carolinian ecoregion, was distinct from all others, implicating the Florida Peninsula as a phylogeographic barrier. Biophysical particle-tracking simulations independently recovered the same spatial structure, with modeled larval exchange consistent with the genomic signal across the bank network. These findings reveal that *S. exserta* persists across the northern WTNWA province through a dual reproductive strategy of clonality at the bank scale and episodic larval exchange at the network scale, with implications for the conservation and management of mesophotic octocoral metacommunities.

## 1. INTRODUCTION

### 1.1 Background and importance

Population connectivity, the exchange of individuals (especially larvae) among geographically separated subpopulations, is widely recognized as key to the persistence and resilience of coral species (Jones et al., 2009). Connectivity maintains genetic diversity within populations and facilitates recolonization after disturbances, thereby bolstering the adaptability and recovery potential of coral communities (Baums, 2008; Hock et al., 2017). In ecosystem management and restoration, understanding connectivity is crucial; for example, knowledge of larval dispersal routes helps identify source and sink populations and informs the design of marine protected areas and restoration strategies (Cowen & Sponaugle, 2009; Green et al., 2015; Sale et al., 2010). In short, well-connected coral populations are better able to exchange genetic material and replenish damaged sites, increasing the odds of long-term survival.

Mesophotic coral ecosystems (MCEs), typically found at ∼30–150 m depth, are increasingly recognized as diverse and ecologically important habitats (S. Kahng et al., 2017; Lesser et al., 2018). In the Warm Temperate Northwest Atlantic coastal and shelf (WTNWA) biogeographic province (Spalding et al., 2007), MCEs occur on rocky outcrop reefs, or banks, that host abundant azooxanthellate corals (gorgonians and black corals) and associated fauna (Sammarco et al., 2016). These deeper ecosystems provide many of the same ecosystem functions as shallow coral reefs. They serve as essential fish habitat and support biodiversity, but are distinguished by slower-growing coral communities adapted to low light and high nutrient input from surface waters (Holstein et al., 2019; S. E. Kahng et al., 2010). MCEs are also highly susceptible to environmental disturbances. For example, the 2010 *Deepwater Horizon* (DWH) oil spill exposed several mesophotic reefs in the Northern Gulf ecoregion to oil for over a month, resulting in significant coral injury. Post-spill surveys at impacted sites found that roughly one-third of large gorgonian colonies showed tissue damage, broken branches, and other signs of stress (Etnoyer et al., 2016; Silva et al., 2016). This event starkly demonstrated that mesophotic reefs, despite their depth, are not refuges from anthropogenic impacts, as pollutants and other stressors can readily reach these communities.

The octocoral *Swiftia exserta* (Ellis & Solander, 1786) is an integral component of mesophotic reefs along the eastern United States from Texas to Florida and up to the Carolinas, spanning the WTNWA, the Tropical Northwestern Atlantic, the North Brazil Shelf and the Tropical Southwestern Atlantic provinces, where it features prominently as a habitat-forming species on hard bottom habitats (Goldberg, 2001). This bright red-orange gorgonian occurs from relatively shallow depths (∼18 m) and commonly down to 200 m, although there are deeper records that need further assessment (Intergovernmental Oceanographic Commission of UNESCO, n.d.; Johnstone et al., 2025). Like other gorgonians, *S. exserta* adds three-dimensional complexity to the benthic habitat as they create shelter and attachment surfaces that support diverse animal assemblages (**Figure 1A**.; (Sánchez et al., 2019)). As an azooxanthellate species reliant on plankton and particulate matter from the water column, *S. exserta* can thrive in low-light conditions (Lange et al., 2025), but remains vulnerable to water-borne pollutants (Frometa et al., 2017). Little was historically known about the life history of *S. exserta*, although recent observations have begun to shed light on its reproductive biology. *S. exserta* appears to be gonochoric (separate sexes) and a broadcast spawner (Johnstone et al., 2025). Fragments maintained in captivity spawned ripe gametes, with fertilized eggs developing into swimming planula larvae within ∼3 days post-spawning and larvae settling by ∼14 days (Johnstone et al., 2025). Larvae are not obligately demersal, as unsettled *S. exserta* larvae remained capable of swimming in the water column for up to two months when no suitable substrate was available (McCauley et al., 2026). This reproductive mode (broadcast spawning with pelagic larvae, with the capacity to extend the pelagic phase in the absence of substrate) suggests *S. exserta* has the physiological capacity for dispersal across considerable distances, potentially linking remote mesophotic populations via larval connectivity.

**FIGURE 1.**
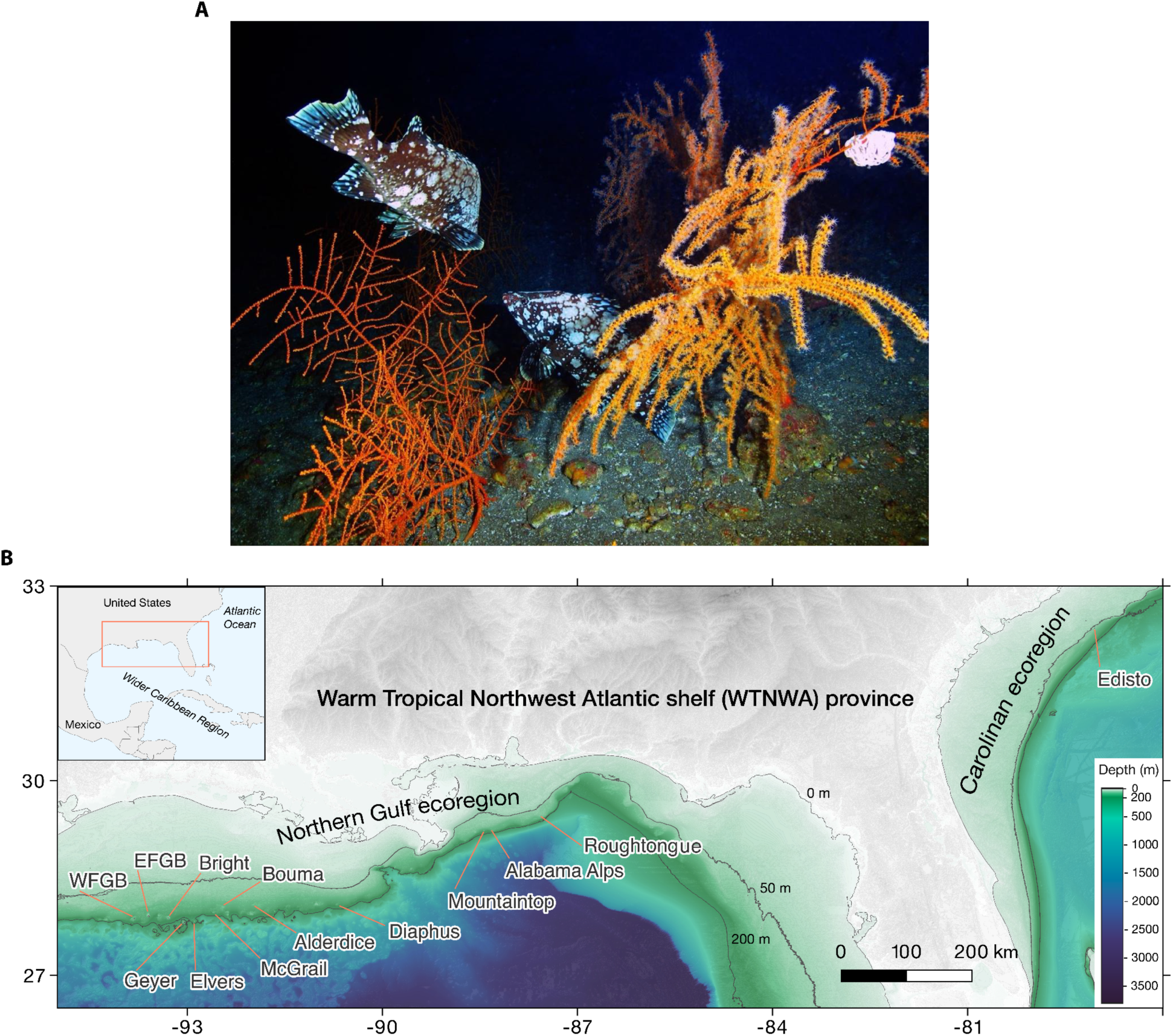
*Swiftia exserta* and sampling locations in the Warm Temperate Northwestern Atlantic shelf (WTNWA) biogeographic province. (A) *S. exserta* colony at Elvers Bank, Northern Gulf ecoregion, (102 m, September 2018). Fish, like the marbled grouper (*Dermatolepis inermis*), and invertebrates, like basket stars are commonly seen near or on this coral. (B) Sampling locations of *S. exserta* across the WTNWA biogeographic province, spanning the Northern Gulf and Carolinian ecoregions. Site abbreviations used in subsequent figures and tables: WFGB = West Flower Garden Banks, EFGB = East Flower Garden Banks, MTR = Mountaintop Reef, AAR = Alabama Alps Reef, RTR = Roughtongue Reef.

Notably, *Swiftia exserta* was among the coral species significantly impacted by the DWH oil spill. Surveys conducted after the spill documented a marked decline in the condition of *S. exserta* colonies at affected mesophotic reefs, with observations of extensive tissue loss, bare skeletons, and abnormal growths on colonies exposed to the oil (Etnoyer et al., 2016; Silva et al., 2016). Laboratory toxicity experiments confirmed that exposure to crude oil and chemical dispersant is lethal to *S. exserta* larvae and polyps (Frometa et al., 2017). Given these impacts, *S. exserta* has been identified as a priority for active restoration efforts in the WTNWA Northern Gulf (Frometa & Etnoyer, 2023). However, the success of these restoration efforts hinges on a deeper understanding of *S. exserta* biology, particularly its population genetic structure and connectivity, to ensure that restored populations are genetically diverse, self-sustaining, and sourced from appropriate stocks.

### 1.2 Knowledge gaps

Despite their ecological importance, the genetic structure and dispersal patterns of mesophotic corals like *Swiftia exserta* remain poorly understood. Most research on coral population genetics and connectivity has focused on shallow-water reef-building corals as reviewed by (Banha et al., 2025; Eyal et al., 2021; Laverick & Rogers, 2020), whereas mesophotic and deep-sea corals have received comparatively little attention. Logistical challenges, including the difficulty of accessing deep reefs and obtaining sufficient samples, have limited studies in these environments (Brooke et al., 2023; Turner et al., 2017). While mesophotic stony coral connectivity has received growing attention (Bongaerts et al., 2017; Studivan & Voss, 2018b), fundamental questions about how mesophotic octocoral populations are connected (or isolated) over tens to hundreds of kilometers are largely unanswered. The paucity of data on larval dispersal and gene flow in deeper coral ecosystems is a critical knowledge gap that impedes our ability to predict their recovery potential and to design effective conservation measures (Cowen & Sponaugle, 2009; Holstein et al., 2019). In the case of *S. exserta*, previous surveys have established where the species occurs and documented its post-spill decline (Etnoyer et al., 2016; Goldberg, 2001; Silva et al., 2016), but we still lack clarity on whether populations across the Northern Gulf function as a single genetic stock or as distinct subpopulations with limited exchange.

Whether populations exchange migrants depends in large part on how larvae are produced and dispersed. Reproductive mode sets the potential scale of larval dispersal in corals, with brooded larvae recruiting close to their parents and broadcast-spawned larvae capable of traveling much farther (Coelho & Lasker, 2016; Harrison & Wallace, 1990). How much of that potential is realized depends on temperature-dependent development, larval behavior, and the timing of release relative to local circulation, with realized dispersal distances often correlating poorly with planktonic duration across marine taxa (Burgess et al., 2016; Shanks, 2009). Whether *S. exserta*’s broadcast spawning translates into extensive connectivity or is offset by retentive circulation and larval behavior remains unresolved.

The two reproductive modes available to octocorals differ in how far offspring travel, and whether offspring are genetically novel at all. Sexual reproduction generates genotypic diversity through recombination, increasing the probability that at least some fraction of a population will carry genotypes capable of tolerating novel or intensifying stressors (Baums, 2008). Cloning trades that diversity for demographic reliability: in the Caribbean octocoral *Plexaura kuna*, fragment-derived colonies were estimated to have up to 25-fold greater probability of reaching reproductive size than larval recruits (Lasker, 1990). Because disturbance both selects for genotypic diversity and physically generates fragments, clonality has been found to increase with disturbance in some corals (Coffroth & Lasker, 1998; Foster et al., 2013).

Clonality also complicates the interpretation of population genetic data in ways that other reproductive traits do not. Asexual reproduction through fragmentation or colony fission produces ramets genetically identical to the original individual, decoupling colony abundance from genotypic abundance, in which a reef may support many colonies but few distinct genets (Lasker, 1990; McFadden, 1991, 1997). The consequences may be greatest in habitats that are patchy and isolated, such as mesophotic banks, where hard substratum is discontinuous, disturbance is episodic, and larval supply is temporally variable. Once a genotype has been established under such conditions, ramet production allows a founder to occupy locally available space without requiring further successful settlement (McFadden, 1991, 1997). However, for mesophotic octocorals like *S. exserta*, the extent of clonal reproduction and its contribution to population connectivity are virtually unknown. Without that baseline, genotypic and colony-level patterns cannot be reliably distinguished.

### 1.3 Study objectives

Considering the above knowledge gaps, the present study aims to resolve the population genetic structure and connectivity of *Swiftia exserta* in the WTNWA coastal and shelf biogeographic province. We took a population genomics and biophysical modeling approach, utilizing genome-wide SNP markers (derived from Restriction-Associated DNA Sequencing, RADseq) to achieve high resolution detection of genetic differentiation among *S. exserta* colonies from multiple mesophotic reef sites. The first objective is to quantify the genetic structure of *S. exserta* populations – identifying discrete genetic stocks (if they exist) and assessing levels of genetic diversity within and between sites. The second objective is to characterize patterns of connectivity among these populations, inferring the degree of larval exchange and directional dispersal across the region. We hypothesize that *S. exserta* will exhibit measurable spatial genetic structure across the sampled range, reflecting the species’ mesophotic habitat patchiness and larval dispersal capabilities.

## 2. MATERIALS AND METHODS

### 2.1 Sample collection and preservation

*Swiftia exserta* colonies were sampled from 13 sites in the WTNWA province, 12 in the Northern Gulf ecoregion and one in the Carolinian ecoregion, at depths between 50 and 100 meters (**Figure 1B**). The 2010 *Deepwater Horizon* oil spill directly impacted *S. exserta* populations at two of these sites in the Pinnacles Trend area (Alabama Alps (AAR) and Rough Tongue Reef (RTR)) (Etnoyer et al., 2016), Mountaintop Reef was not included in the oil spill assessment but was included as a sampling site here. Collections took place during expeditions in 2017 (OP17 MSV *Ocean Project* & RV *Manta*, ROV *Comanche* & ROV *Mohawk*), 2018 (MT18 RV *Manta*, ROV *Mohawk*), and 2019 (PE1925 RV *Pelican*, ROV *Global Explorer*). Individual coral colonies were imaged before and after tissue collection. Tissue samples were collected by removing a small distal branch using hydraulic manipulators mounted on remotely operated vehicles. Samples were then stored in insulated containers until recovery of the vehicles by the surface vessel. Subsamples of each specimen were preserved in liquid nitrogen or 95% ethanol and stored at −80 °C.

### 2.2 DNA extraction and library preparation

To characterize the genetic diversity of *S. exserta*, we performed a reduced representation DNA sequencing (i.e., RAD-seq) (Baird et al., 2008; Reitzel et al., 2013). DNA was purified using the salting-out protocol for extracting high-molecular weight DNA from octocorals (Herrera, 2022). DNA was subsequently cleaned using the Qiagen DNeasy PowerClean Pro Clean Up Kit, and DNA integrity and purity were assessed by visual inspection on a 1% agarose gel and a Nanodrop spectrophotometer (Nanodrop Technologies), respectively. DNA concentration was determined and normalized using a Qubit 4.0 fluorometer with broad-range reagents (Invitrogen). Species identification was confirmed through DNA barcoding of the *mtMutS* mitochondrial gene as described by (Frometa et al., 2021). Floragenex Inc (Eugene, OR) performed RAD sequencing library preparation utilizing the 6-cutter PstI restriction enzyme on quality-checked and concentration-normalized high-molecular-weight DNA. Using the program PredRAD (Herrera et al., 2015), we predicted 75,550 cleavage sites in coral genomes with the PstI restriction enzyme. Libraries were dual-barcoded and sequenced on an Illumina Hi-Seq 4000 platform using single-end bp reads (1×100).

### 2.3 Reference genome assembly for *Swiftia exserta*

A *de novo* reference genome assembly was generated for *S. exserta* to support read mapping and variant discovery. The reference specimen, SH10214, was collected in the Northern Gulf at Roughtongue Reef during expedition OP17 aboard the MSV *Ocean Project*. The sample was collected by ROV *Comanche* during dive CM14 at station 37-4 on 27 July 2017 at 11:57 from 65 m depth at 29.43857°N, 87.57548°W. The specimen was flash-frozen in liquid nitrogen immediately after collection and stored at −80 °C until DNA extraction. High molecular weight genomic DNA was extracted from frozen tissue and purified using the protocol described above. PacBio continuous long-read sequencing generated 67.2 Gb from one SMRT Cell, comprising 5,040,439 polymerase reads and 67,229,257,044 polymerase-read bases. For polymerase reads longer than 8 kb, the mean read length was 13,338 bp and the polymerase read N50 was 26,896 bp.

The initial assembly was generated from PacBio continuous long reads using FALCON (Chin et al., 2016) and polished with Arrow. The polished assembly was scaffolded with Omni-C proximity ligation data using HiRise, which uses long-range linkage information to identify misjoins, score candidate joins, and build chromosome-scale scaffolds (Putnam et al., 2016). For HiRise scaffolding, Omni-C read pairs were mapped to the draft assembly with the SNAP read mapper (Zaharia et al., 2011). HiRise estimated a likelihood model relating contact frequency to genomic distance, then used this model iteratively to break putative misjoins and join scaffolds exceeding the pipeline-specific likelihood threshold. Assembly statistics were calculated with QUAST v5.2.0 (Gurevich et al., 2013). Population genomic reads were mapped to the final full HiRise assembly, and the 17 largest scaffolds were retained as a putative chromosome-scale subset for analyses requiring chromosome-scale coordinates.

### 2.4 Sequence quality control and snp discovery

Raw sequence RADseq reads were de-multiplexed and quality filtered using the process_radtags program in Stacks v2.54 (Catchen et al., 2013) with the following flags: -- inline_null, -r, -c, and -q, with default values. BWA v0.7.17 (Li & Durbin, 2009) was used to map reads to the *S. exserta* chromosome-level assembly. Single-nucleotide polymorphisms (SNPs) were called from mapped reads using the Stacks ref_map.pl pipeline. Assembled loci were then processed in the populations program of Stacks to export a SNP matrix in vcf format for downstream analyses.

### 2.5 SNP and individual filtering

A two-step filtering approach was applied to ensure data quality. First, loci with >10% missing data and individuals with >30% missing data were removed. As all loci met the missing data criterion, only individual-level filtering was necessary. Loci with unusually high observed heterozygosity (Ho > 0.5), potentially indicative of paralogous sequences or genotyping errors, were identified and removed (n = 304). BayeScan v2.1 was used to identify loci potentially under selection in *Swiftia* populations by analyzing SNP loci across 13 populations (Foll & Gaggiotti, 2008). BayeScan implements a Bayesian method that directly estimates the probability of each locus being under selection by decomposing F_ST_ coefficients into population-specific and locus-specific components. This analysis was performed using the default parameters of 20 pilot runs, 50,000 iterations for burn-in, followed by 5,000 samples with a thinning interval of 10. Initial detection yielded 194 loci under positive, diversifying selection (positive alpha values, log10(PO) >2). To minimize false positives, only loci meeting both global F_ST_ > 0.35 and q-value < 0.05 were retained as outliers and subsequently removed from the neutral dataset. This conservative threshold identified 18 high-confidence outlier loci showing strong differentiation with F_ST_ values ranging between 0.35-0.49, which is substantially higher than the global genome-wide average of 0.112 (**Figure S1**).

### 2.6 Clone detection and spatial distribution

Clones were identified using the poppr package in R (Kamvar et al., 2014). The distribution of all pairwise genetic distances was examined across all populations to determine the appropriate genetic distance threshold for clone detection. This revealed a bimodal distribution, indicating distinct clusters representing clonal relationships versus genetically distinct individuals. Based on this distribution, we applied a conservative threshold of 2,000 allelic differences using the *mlg.filter* function to collapse samples into unique multilocus genotypes (MLGs) for downstream population genetics analysis.

We define a genet as a genetically distinct individual and a ramet as each physical colony sharing an identical multilocus genotype with that genet; a genet represented by two or more ramets is referred to as a clonal genet. The spatial distribution of clonal genets was analyzed using geographic coordinates and depth. For each clonal genet, the mean and maximum geographic distances between clones were calculated using the Haversine formula through the *distm* function from the geosphere package (Hijmans, 2024). Clonal genets were validated through multiple approaches, examining genetic distances and detailed genotype comparisons within each genet. Initial validation compared genetic distances within genets to typical within-site versus between-site genetic distances, confirming that legitimate clones showed consistently low genetic distances (<2,000 allelic differences). Cross-site genets were further investigated using locus-by-locus genotype comparisons to identify the specific number and positions of SNP differences between putative ramets. The final clone-corrected dataset was created by retaining one representative individual per validated MLG per sampling site using the *clonecorrect* function with population stratification.

Loci with unusually high observed heterozygosity (Ho > 0.5) that might indicate paralogous sequences or genotyping errors were identified and removed (n = 304). Loci significantly deviating from Hardy-Weinberg Equilibrium (HWE) were identified using the *hw.test* function from the pegas package (Paradis, 2010), with 1,000 permutations, and loci with p < 0.01 were excluded. The final clone-corrected, neutral dataset consisted of 192 individuals representing 189 unique multilocus genotypes across 13 populations, with the difference reflecting retention of some genotypes across multiple populations. Loci that significantly deviated from HWE expectations (p < 0.01) were removed to create the final clone-corrected, neutral dataset for downstream analyses. Following removal of 18 high-confidence outlier loci identified by BayeScan, the final filtered dataset consisted of 16,746 loci with markers adhering to neutral evolutionary expectations (**Table 1**).

**Table 1.** Filtering pipeline for the *Swiftia exserta* neutral SNP dataset. For each processing step, the number of individuals, loci, and populations retained is reported.

| Step | Description | Individuals | Loci | Populations |
| --- | --- | --- | --- | --- |
| Raw dataset | Post loci >10% missing / individuals >30% missing filtering | 288 | 23869 | 13 |
| Clone correction | *clonecorrect()* retained one representative per validated MLG per population | 192 | 23869 | 13 |
| Ho filtering | Loci with observed heterozygosity $H_o > 0.5$ removed | 192 | 23565 | 13 |
| HWE filtering | Loci deviating from HWE at $p < 0.01$ removed (hw.test, 1000 permutations, seeded) | 192 | 16764 | 13 |
| Outlier removal (BayeScan) | 18 high-confidence outlier loci removed ($F_{ST} > 0.35$, $q < 0.05$) | 192 | 16746 | 13 |
| Final neutral dataset | Clone-corrected, neutral dataset used for downstream analyses | 192 | 16746 | 13 |
| Northern Gulf subset | Gulf populations with $n \geq 10$ individuals; Edisto excluded | 158 | 16746 | 9 |
| Northern Gulf subset (LD-pruned, ADMIXTURE) | Linkage disequilibrium-pruned used only for ADMIXTURE | 158 | 5419 | 9 |
| Gulf-Carolinian subset | Gulf populations with $n \geq 10$ individuals, Edisto included | 174 | 16746 | 10 |
| Gulf-Carolinian subset (LD-pruned, ADMIXTURE) | Linkage disequilibrium-pruned used only for ADMIXTURE | 174 | 5629 | 10 |

### 2.7 Site-specific genetic diversity

Site-specific genetic diversity was quantified for the final clone-corrected, neutral SNP dataset. Observed heterozygosity (Ho) and gene diversity (He, corrected for sample size; hereafter Hs) were calculated per locus per population using the *basic.stats* function in the hierfstat package, and averaged across all loci to obtain site-level estimates (Goudet, 2005). The inbreeding coefficient (FIS) was calculated at 1-(Ho/Hs) per locus and averaged per site. Allelic richness was calculated using *allelic.richness* in hierfstat, rarefied to the smallest population sample size (n = 5, Western Flower Garden Banks (WFGB)) to allow unbiased comparison across sites of unequal sample size. The percentage of polymorphic loci per site was calculated as the proportion of loci with more than one observed allele count among genotypes individuals at that site. Private alleles were defined as alleles present in only one population and absent from all others, calculated at the allele level for each biallelic SNP. Site-level summary statistics were reported for sample sizes at each stage of the filtering pipeline (raw, post-clone correction, and in the final analyzed dataset) and the number of validated within-site genets identified for each population.

### 2.8 Potential connectivity

Ocean conditions from September to November of 2014 and 2015 were simulated using the Coastal and Regional Ocean Community (CROCO) model in its hydrostatic configuration (Auclair et al., 2019). The CROCO domain covered 24°N to 31°N and 98°W to 82°W at 1 km horizontal resolution, and with 50 terrain-following vertical levels, providing enhanced resolution near the ocean floor. The model was forced at the surface using heat and momentum fluxes from the Navy Global Environmental Model (NAVGEM) and nudged at the boundaries to the Hybrid Coordinate Ocean Model – Navy Coupled Ocean Data Assimilation (HYCOM-NCODA) analysis system (Cummings, 2005; Cummings & Smedstad, 2013). Daily freshwater input from the ten major rivers within the domain (Sun et al., 2022), and the dominant tidal constituents (Egbert & Erofeeva, 2002) were included. The model bathymetry was generated using ETOPO2 (National Geophysical Data Center/NESDIS/NOAA/U.S. Department of Commerce, 2001), a 2-minute Gridded Global Relief topography dataset. This model set-up has been previously validated and used to assess the dispersal of Red Snapper larvae (Zhou et al., 2024) and the connectivity potential between mesophotic areas in the Flower Garden Banks Marine Sanctuary (Lopera et al., 2025) and (Pittoors et al., 2025).

Ichthyop v3.3.16 was used to simulate the three-dimensional Lagrangian particle dispersal during the *S. exserta* spawning seasons of September and October (Lett et al., 2008) in 2014 and 2015. The years were chosen because they represented climatological currents (2014) and high eddy kinetic energy with an extended Loop Current for part of the year and a Loop Eddy moving east to west across the domain of interest (2015). Therefore, 2014 is considered a year with “normal” conditions, whereas 2015 represents a more “extreme” scenario. On September 25 and October 5 of each year, 90,000 particles were released 50 cm above the seafloor to avoid being impacted by the model no-flow conditions at the bottom given the vertical discretization within ∼5 km² areas (again accounting for model discretization) surrounding the sampling sites and advected for a period of 60 days. This duration reflects the documented planktonic larval duration of *S.* exserta: under substrate-limited laboratory conditions, larvae survived in the water column for up to two months prior to settlement (McCauley et al., 2026). Based on observations of *S. exserta* biology published in (Johnstone et al., 2025), particles were assumed to be neutrally buoyant for the first day following broadcast spawning, followed by three days during which they were positively buoyant and confined at or close to the ocean surface. Afterward, they became negatively buoyant and remained near the ocean bottom for the rest of their duration. Particles that descended to depths of 500 m or greater were considered unsuccessful settlers and removed from the simulation, as suitable hard substrate habitat for *S. exserta* does not occur at these depths and larval survival under associated pressure and temperature conditions is not expected.

### 2.9 Population structure and genetic differentiation

All population-level analyses were performed on two datasets derived from the neutral dataset. Four Northern Gulf sites with fewer than 10 genets following clone removal (WFGB, Geyer, Bright, and RTR) were excluded to ensure robust parameter estimation. The Northern Gulf dataset (GoM10) consisted of 158 individuals across the remaining 9 Gulf populations. The Gulf-Carolinian dataset included 174 individuals across 10 populations, extending the Northern Gulf dataset to include Edisto off South Carolina. Pairwise genetic differentiation among sampling sites was estimated using the Weir and Cockerham (1984) F_ST_ estimator implemented in SNPrelate (Zheng et al., 2012). To test for isolation by distance (IBD), Mantel tests were conducted using the *mantel* function in the vegan package (Oksanen, 2010; Ter Braak C Weedon J, 2026), testing the correlation between linearized genetic differentiation [F_ST_/(1-F_ST_)] and the natural log of geographic distance (km) calculated from site coordinates using the Haversine formula. Statistical significance was assessed using 9,999 permutations. To evaluate depth as an alternative or confounding driver of observed structure, linearized genetic differentiation was additionally tested against pairwise depth distance (absolute difference in mean depth, m), and a partial Mantel test evaluated genetic differentiation against geographic distance while controlling for depth.

Analysis of molecular variance (AMOVA; (Excoffier et al., 1992)) was conducted using the *poppr.amova()* function in the R package poppr, with Euclidean distances calculated on mean-imputed allele frequencies. For the Northern Gulf dataset, a single-level AMOVA partitioned molecular variance within and across populations. For the Gulf-Carolinian dataset, both a single-level AMOVA and a two-level hierarchical AMOVA were performed; the hierarchical analysis partitioned variance among ecoregions (Northern Gulf vs. Carolinian), across populations within regions, and within population. Statistical significance of Φ-statistics was assessed via permutation tests (9,999 permutations).

Prior to population structure analysis, linkage disequilibrium (LD) pruning was performed using SNPRelate to remove highly correlated SNPs, applying a sliding window of 500 kb, an LD threshold of |r| ≤ 0.1, a minor allele frequency filter of 0.05, and a maximum missing data rate of 0.1. (Zheng et al., 2012). Population structure was investigated using three complementary approaches: Discriminant Analysis of Principal Components (DAPC), Uniform Manifold Approximation and Projection (UMAP), and admixture analysis, and haplotype-based coancestry analysis. All analyses were performed on the neutral dataset containing 192 individuals from 13 populations after clone correction and filtering, unless otherwise specified.

Discriminant Analysis of Principal Components (DAPC) was performed using the adegenet package v2.1.10 in R (Jombart et al., 2010; Jombart & Ahmed, 2011). Two approaches were employed: (1) clustering without prior population assignment to identify optimal genetic clusters (K), and (2) analysis with geographic priors using predefined sampling locations. For the clustering approach without priors, the optimal number of principal components was determined using the *optim.a.score()* function, which identified 19 components as optimal for balancing information retention and overfitting avoidance. The optimal number of genetic clusters was determined using Bayesian Information Criterion (BIC) through the *find.clusters()* function, evaluating K values from 1 to 20. For DAPC with geographic priors, populations were defined by sampling location, and the analysis retained 19 principal components and 6 discriminant functions. The percentage of variance explained by each discriminant axis was calculated, and correlations between discriminant function scores and geographic variables (longitude, latitude, depth) were assessed using Spearman rank correlation tests with significance set to α = 0.05. Uniform Manifold Approximation and Projection (UMAP) was applied to DAPC scores to provide an additional dimensionality reduction approach that preserves both local and global data structure, implemented using the *uwot* package v0.2.10.0 in R (Konopka, 2023).

The LD-pruned dataset was converted to PLINK binary format using PLINK v1.9 (Purcell et al., 2007) for input to ADMIXTURE. Admixture analysis was conducted using ADMIXTURE v1.3.0 to estimate individual ancestry proportions (Alexander et al., 2009). ADMIXTURE was run for K values ranging from 1 to 10, with 10 replicate runs per K value to assess consistency across random starting seeds. The optimal K value was determined based on cross-validation (CV) error, evaluated across the Northern Gulf, and Gulf-Carolinian datasets.

Haplotype-based co-ancestry among individuals was estimated using fineRADstructure (Malinsky et al., 2018). RADpainter was first used to convert phased RAD haplotypes from the VCF to a haplotype input file, after which loci were reordered by linkage disequilibrium using a random sample of 500 loci. RADpainter *paint* was then used to compute a pairwise co-ancestry matrix from the reordered haplotype data. The resulting coancestry matrix was analyzed in fineStructure, which employs a Markov chain Monte Carlo (MCMC) algorithm to infer population clustering from haplotype coancestry without requiring predefined population assignments. The MCMC was run for 100,000 burn-in iterations followed by 100,000 sampling iterations with a thinning interval of 1,000, retaining 100 posterior samples. A population tree was subsequently inferred using 10,000 additional tree-search iterations. Analyses were performed independently on the Northern Gulf and Gulf-Carolinian datasets. Population-averaged coancestry matrices were computed by averaging individual-level coancestry values within and between sampling sites visualized as heatmaps with populations ordered west to east.

## 3. RESULTS

### 3.1 De novo reference genome assembly

The final HiRise-scaffolded *S.exserta* assembly contained 1,063 scaffolds totaling 428,812,460 bp (157x coverage). Scaffold lengths ranged from 1,179 bp to 56,230,999 bp. GC content was 36.80%, and gap content was low at 20.89 ambiguous bases per 100 kbp. The assembly had a scaffold N50 of 18,534,683 bp, N90 of 233,827 bp, auN of 28,317,658.9 bp, L50 of 6 scaffolds, and L90 of 57 scaffolds.

The 17 largest scaffolds were interpreted as putative chromosome-scale scaffolds. These scaffolds totaled 373,046,731 bp, representing approximately 87.0% of the full assembly. Within this subset, scaffold lengths ranged from 6,398,999 bp to 56,230,999 bp. GC content was 36.80%, and gap content was 23.03 ambiguous bases per 100 kbp. All population genetic analyses were performed on genotype data from reads mapped to the putative assembled chromosomes.

### 3.2 Clone detection and spatial distribution

Pairwise genetic distances among all 288 sampled individuals revealed a bimodal distribution within sampling sites, indicating the presence of clonal genotypes alongside genetically distinct individuals (**Figure 2A**). Applying a conservative threshold of 2,000 allelic differences and validation of putative cross-site genets, collapsed the dataset into a final clone-corrected set of 192 individuals across 13 populations, with no genet spanning multiple sites. Following removal of one representative per validated MLG per site via *clonecorrect*, 115 (40%) individuals belonged to one of the 28 validated genets. Within these genets, pairwise genetic distances average 847 allelic differences (range: 152-1,986), well below the applied threshold. Pairwise genetic distances among non-clonal individuals in the clone-corrected dataset averaged 6,476 allelic differences (range: 2,936-8,019), confirming separation between clonal and non-clonal relationships.

**FIGURE 2.**
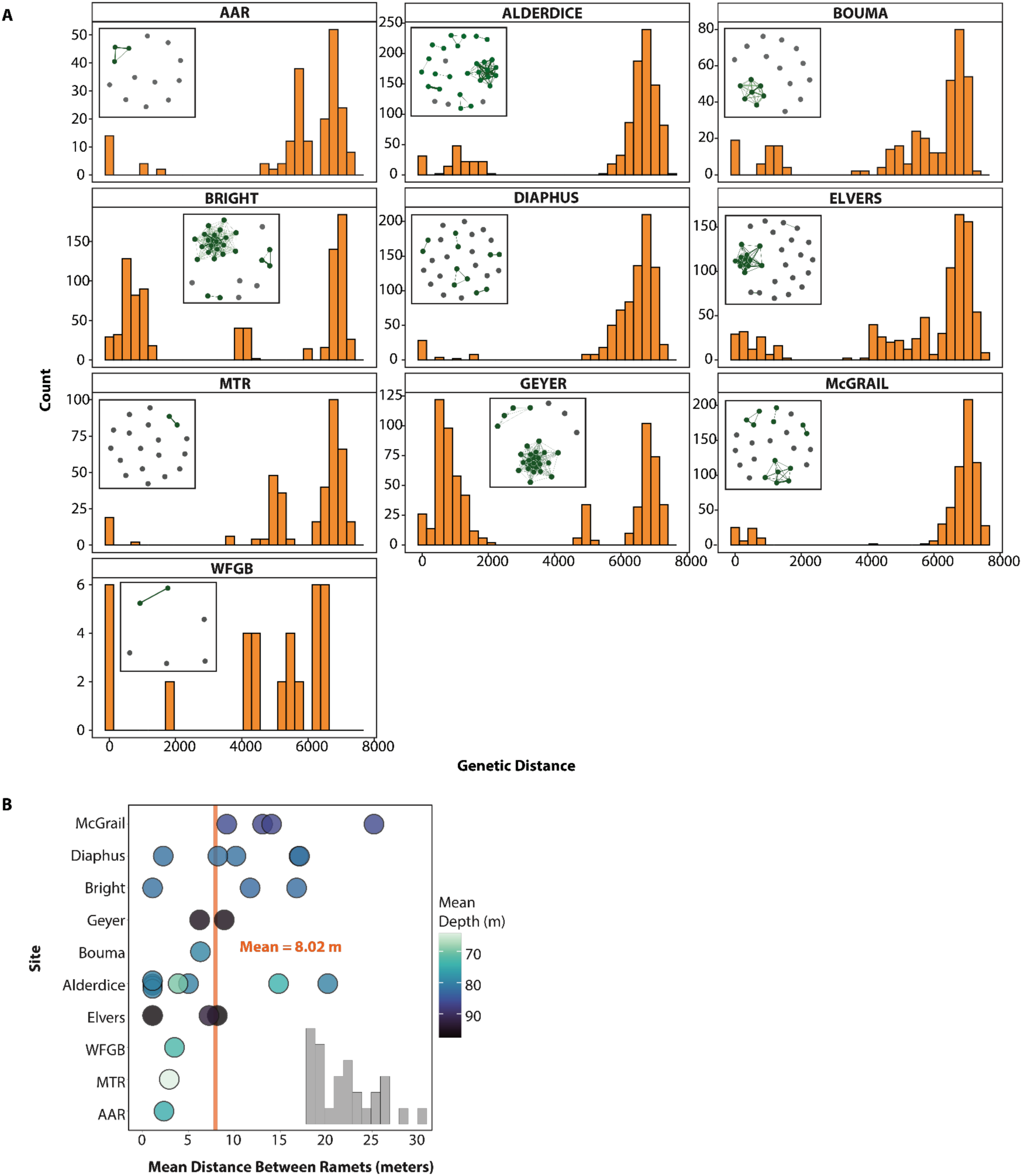
Clonal structure and spatial distribution of *Swiftia exserta* colonies across the WTNWA Northern Gulf ecoregion populations. (A) Frequency distributions of pairwise genetic distances (number of allelic differences between individuals) among colonies within each sampling site. Inset network diagrams depict genet membership within each site, where nodes represent individual colonies and green edges connect members of the same genet (grey nodes indicate non-clonal individuals). (B) Mean geographic distance (meters) between ramets of the same genet at each sampling site. The grey inset histogram shows the frequency distribution of mean clone distances across all sites. Panels are shown only for sites with at least one detected clonal genet; EFGB, RTR, and Edisto are omitted, as no clonal genets were detected at these sites.

Genets exhibited a range of spatial extents, with mean distances between ramets ranging from 0 to 29.8 m (overall mean: 8.02 m; median: 7.7 m) (**Figure 2B**). Genet size ranged from 2 to 19 ramets. Individual genet mean depths ranged from 64.2 m (MTR) to 96.0 m (Geyer). Sites harboring the greatest number of clonal genets were Alderdice ( n = 8), Diaphus (n = 5), and McGrail (n = 4). No clonal genets were detected at East Flower Garden Banks (EFGB), RTR, or Edisto.

### 3.3 Site-specific genetic diversity

Site-specific genetic diversity was broadly consistent across the sampling range (Table 2). Observed heterozygosity ranged from 0.088 (WFGB) to 0.148 (Geyer), and the percentage of polymorphic loci ranged from 33.4% (WFGB) to 81% (RTR), likely reflecting differences in sample size (N = 5-26). Edisto had the greatest number of private alleles (n = 59), consistent with its genetic distinctness from Gulf of Mexico populations identified by DAPC and ADMIXTURE. FIS values were generally low and near zero across sites (range: −0.003-0.235), with the highest value observed at WFGB, the smallest sampled population (**Table 2**).

**Table 2.** Site-specific sampling and genetic diversity summary for *Swiftia exserta* across all 13 sampled populations. N raw, N clone-corrected, indicate individual counts before clone correction, after clone correction, and in the final neutral dataset, respectively. Ho, observed heterozygosity; He, gene diversity (*Hs*); FIS, inbreeding coefficient. Allelic richness is rarefied to the smallest population sample size (n = 5, WFGB). Percentage of polymorphic loci is calculated as the proportion of loci with more than one observed allele count within each site. Private alleles indicate alleles present in only one population. Sites WFGB, Bright, and Geyer (N < 10) were included in site-level diversity summaries but excluded from population-structure analyses (Northern Gulf dataset) due to insufficient sample size for reliable inference.

| Site | Mean lat. | Mean long. | Min depth (m) | Max depth (m) | N raw | N clone corrected | N clonal genets | Ho | He | FIS | Allelic richness | % polymorphic | No. private |
| --- | --- | --- | --- | --- | --- | --- | --- | --- | --- | --- | --- | --- | --- |
| AAR | 29.2522471 | -88.33845 | 71 | 79 | 14 | 12 | 1 | 0.1219 | 0.1475 | 0.1286 | 1.1462 | 63.4599 | 18 |
| Alderdice | 28.0763961 | -91.983528 | 60 | 81 | 31 | 13 | 8 | 0.1346 | 0.1476 | 0.0599 | 1.1471 | 68.6612 | 28 |
| Bouma | 28.0716529 | -92.467345 | 78.3 | 80.1 | 19 | 13 | 1 | 0.1313 | 0.1446 | 0.0613 | 1.1441 | 66.6308 | 15 |
| Bright | 27.8973293 | -93.327541 | 82 | 83.4 | 29 | 8 | 3 | 0.1443 | 0.1449 | -0.0026 | 1.1448 | 52.8007 | 22 |
| Diaphus | 28.0877807 | -90.703271 | 81 | 100 | 28 | 22 | 5 | 0.1384 | 0.1472 | 0.0426 | 1.1470 | 78.3710 | 2 |
| EFGB | 27.968236 | -93.613004 | 73 | 76 | 20 | 20 | 0 | 0.1100 | 0.1400 | 0.1645 | 1.1391 | 66.7801 | 6 |
| Edisto | 32.366232 | -79.038371 | 47.5 | 65 | 16 | 16 | 0 | 0.1325 | 0.1483 | 0.0824 | 1.1477 | 65.1021 | 59 |
| Elvers | 27.8535916 | -92.922786 | 92.2 | 98.6 | 29 | 18 | 2 | 0.1400 | 0.1471 | 0.0340 | 1.1469 | 73.2593 | 14 |
| MTR | 29.2324137 | -88.438042 | 61 | 64.9 | 19 | 18 | 1 | 0.1389 | 0.1453 | 0.0292 | 1.1451 | 72.8472 | 4 |
| Geyer | 27.8492378 | -93.05788 | 95 | 97 | 26 | 5 | 2 | 0.1485 | 0.1507 | -0.0006 | 1.1504 | 47.7666 | 28 |
| McGrail | 27.9621412 | -92.821518 | 84 | 91.2 | 25 | 16 | 4 | 0.1454 | 0.1514 | 0.0278 | 1.1512 | 74.1729 | 13 |
| RTR | 29.4391669 | -87.575613 | 64 | 68 | 26 | 26 | 0 | 0.1403 | 0.1494 | 0.0512 | 1.1492 | 80.9566 | 3 |
| WFGB | 27.8985925 | -93.814013 | 73 | 79 | 6 | 5 | 1 | 0.0875 | 0.1319 | 0.2350 | 1.1249 | 33.3990 | 3 |

### 3.4 Population genetic structure

Pairwise F_ST_ values among Northern Gulf populations were low, ranging from 0.001 (Elvers-McGrail: Diaphus-MTR) to 0.028 (EFGB-AAR), (**Figure 3A, Supplementary Figure 2A**). EFGB exhibited the highest differentiation from all other Northern Gulf populations (F_ST_ range: 0.019-0.028), while geographically proximate population pairs such as Elvers-McGrail and Diaphus-MTR showed near-zero differentiation (F_ST_ = 0.001). Edisto exhibited substantially elevated differentiation from all Northern Gulf populations (F_ST_ range = 0.031-0.048), with the greatest divergence detected against EFGB (F_ST_ = 0.048) and the least against MTR (F_ST_ = 0.031; **Figure 3B**).

**FIGURE 3.**
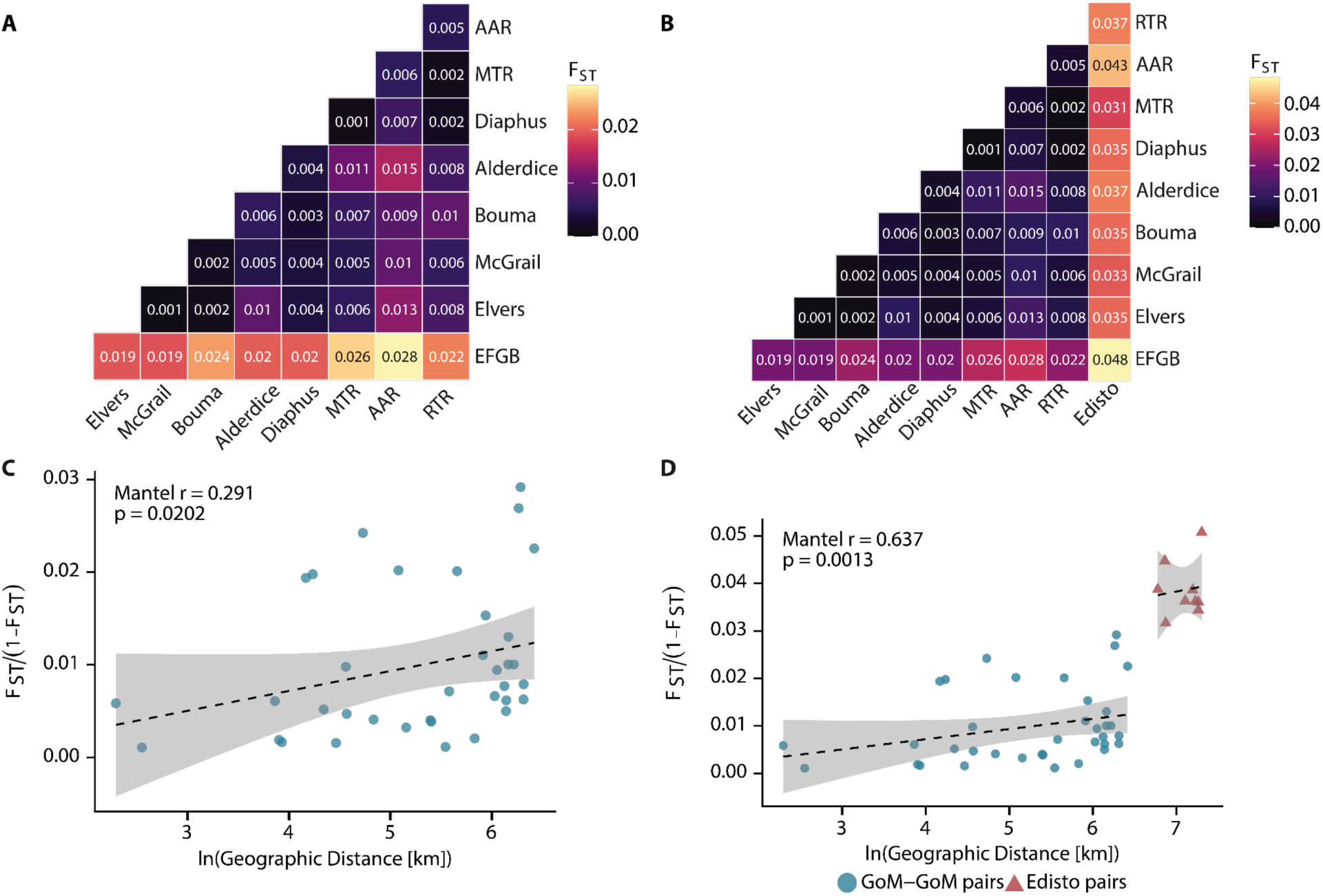
Pairwise F_ST_ and isolation-by-distance (IBD) analyses for *Swiftia exserta* populations. Triangular heatmaps are shown for (A) the Northern Gulf dataset and (B) the Gulf-Carolinian dataset, with populations arranged from west (bottom) to east (top and right).Mantel tests of the relationship between linearized genetic differentiation [F_ST_/(1-F_ST_)] and the natural log of geographic distance (km) are shown for (C) the Northern Gulf dataset and (D) Gulf-Carolinian dataset. Each point represents a pairwise population comparison. Dashed lines indicate the ordinary least squares regression fit with 95% confidence intervals (shaded).

A significant positive relationship between linearized genetic differentiation [F_ST_/(1-F_ST_)] and the natural log of geographic distance was detected among Northern Gulf populations (Mantel r = 0.291, p = 0.020, 9,999 permutations), providing statistical support for isolation by distance as a structuring force within the Gulf (**Figure 3C**). Depth was not significantly correlated with genetic differentiation (Mantel r = −0.208, p = 0.772), and the geographic IBD signal remained significant, becoming marginally stronger, after controlling for depth (partial Mantel r = 0.351, p = 0.002; Table S5). When the Carolinian population, Edisto, was included in the analysis, the IBD signal strengthened considerably (Mantel r = 0.637, p = 0.001; **Figure 3D**), reflecting the pronounced genetic divergence and large geographic distance separating the Atlantic population from all Gulf sites. Depth distance was similarly uncorrelated with genetic differentiation in the Gulf-Carolinian dataset (Mantel r = 0.307, p = 0.145), and the geographic signal persisted after controlling for depth (partial Mantel r = 0.587, p < 0.001; Table S5).

The majority of genetic variation in *S. exserta* was distributed within populations. For the Northern Gulf dataset, 98.7% of variance was attributable to within-population variation, with a small yet statistically significant among-population component (AMOVA; Φ-ST = 0.013, p <0.01). When the Atlantic site Edisto was included (Gulf-Carolinian dataset), among-population variance increases slightly in the single-level analysis (AMOVA; Φ-ST = 0.022, p <0.01) consistent with the addition of a geographically distant sample. The two-level hierarchical AMOVA additionally demonstrated that the Gulf-Atlantic partition accounted for the largest single fraction of molecular variance (4.97%; Φ-CT = 0.050), though this component did not reach statistical significance (p = 0.097), likely reflecting the limited statistical power of a test comparing only two regions was lower in magnitude yet statistically significant (1.18%; Φ-ST = 0.012, p < 0.001), mirroring the significant population structure detected in the Gulf-only analysis.

BIC-based clustering identifies K = 1 as optimal in both the Northern Gulf and Gulf-Carolinian datasets, with BIC increasing monotonically from K = 1 through K = 20 (Gulf: BIC∼K=1∼ = 1,134.4, BIC∼K=2∼ = 1,137.3; Gulf-Carolinian: BIC∼K = 1∼ = 1,251.0, BIC∼K = 2∼ = 1,253.5), indicating no statistical support for discrete genetic clusters in either dataset (**Figure 4A, D**). At K =2, the Gulf-Carolinian dataset resolved a group of 16 individuals corresponding exclusively to Edisto, with all 158 Gulf individuals assigned to a single cluster, confirming the Atlantic-Gulf discontinuity as the dominant structural signal. Within the Gulf, the K = 2 partition separated 25 individuals concentrated at EFGB from the remaining 133, corroborating EFGB’s position as the most genetically differentiated Gulf population across all analyses.

**Figure 4.**
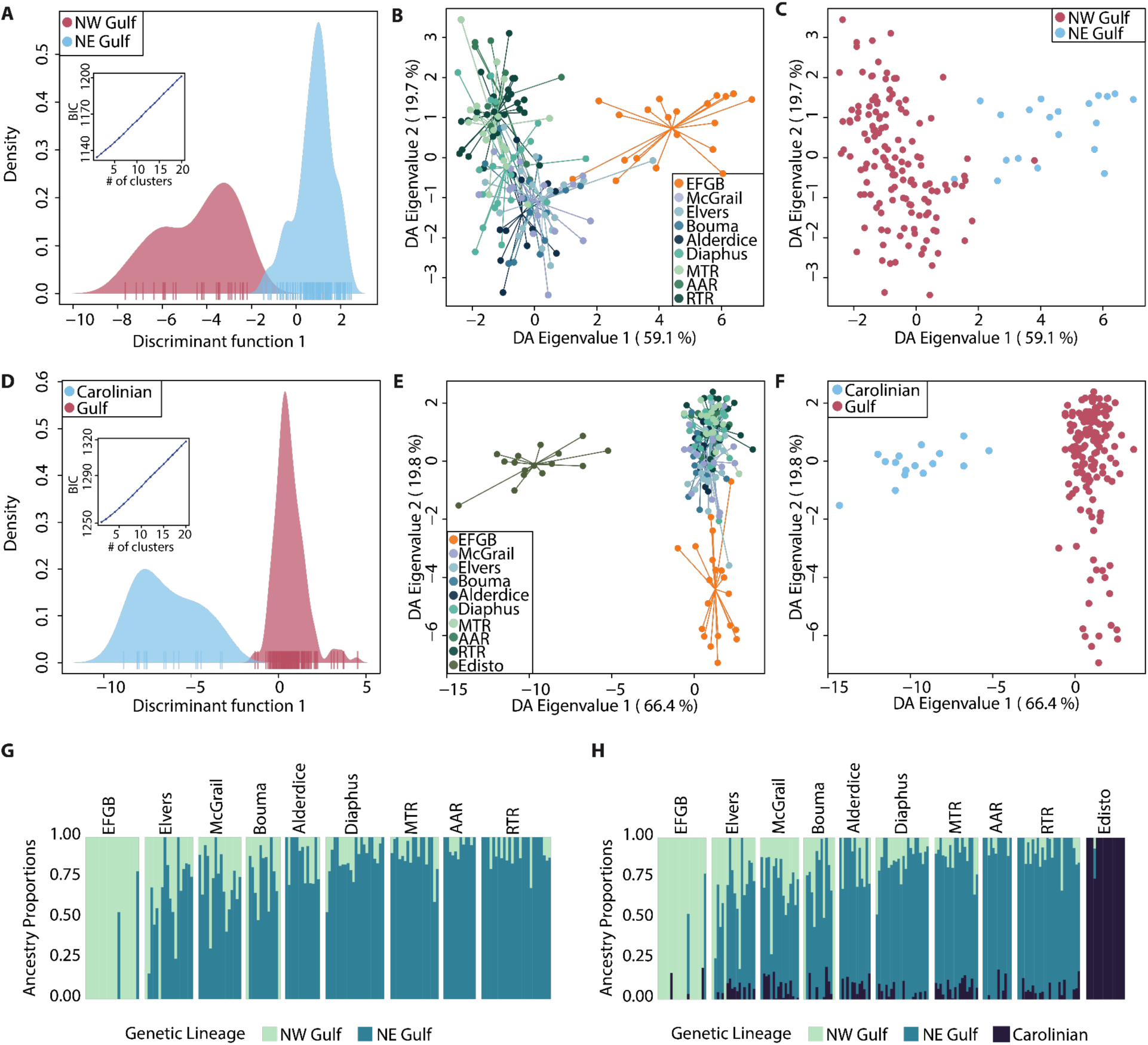
Population Genetic Structure of the octocoral *Swiftia exserta* in the WTNWA province. (A-C) Discriminant Analysis of Principal Components (DAPC) for the Northern Gulf dataset and panels (D-F) show results for the Gulf-Carolinian dataset. (A, D) Density distributions of individual scores along discriminant axis 1 (DA1); insets show Bayesian Information Criterion (BIC) values across K 1-20 clusters. (B,E) Scatter plots of DA eigenvalues 1 and 2 for all individuals with geographic prior assignment, colored by sampling site; lines connect all individuals to their population centroid. (C, F) Same scatter plot as (B,E), with individuals colored by broad genetic grouping. (G,H) ADMIXTURE bar plots (K = 2) for the Northern Gulf and (K = 3) Gulf-Carolinian datasets, respectively, showing individual ancestry proportions arranged west to east. Each vertical band represents an individual; colors indicate proportional membership in each genetic cluster.

DAPC with geographic priors demonstrated a strong geographic signal in genetic variation across both datasets. For the Northern Gulf dataset, DA1 (59.1% of variance) separated EFGB from remaining Gulf populations, while DA2 (19.7% of variance) captured residual geographic variation among central and eastern Gulf sites (**Figure 4B, C**). For Gulf-Carolinian dataset, DA1 (66.4% of variance) separated Edisto from all Gulf populations along a single dominant axis, with DA1 scores showing weak, significant correlations with longitude (ρ = −0.220, p = 0.003), latitude (ρ = −0.195, p = 0.010), and depth (ρ = 0.164, p < 0.031), reflecting the influence of Edisto’s geographic position on this axis (**Figure 4E, F**). DA2 (19.8% of variance) captured the within-Gulf longitudinal gradient, with significant correlations with longitude (ρ = 0.484, p < 0.001) and latitude (ρ = −0.435, p < 0.001), but not depth (ρ = −0.114, p < 0.136). UMAP ordination of DAPC scores supported a continuous gradient of genetic similarity across Gulf populations instead of discrete boundaries, with Edisto isolated from all Gulf populations in multivariate space (**Supplementary Figure 3**).

Admixture analysis identified K = 1 as optimal by cross-validation error in both datasets, with CV error increasing monotonically through K = 10. At K = 2 for the Northern Gulf dataset, we refer to the two resulting clusters as ‘NW Gulf’ and NE ‘Gulf’, reflecting their geographic association with the western (EFGB) and eastern (RTR) ends of the bank network included in this analysis. These notes denote relative positions along a continuous gradient. The NW Gulf-associated component was most prevalent in EFGB individuals but present as a minor component across all Gulf populations, with no discrete geographic boundary (**Figure 4G**). At K = 3 for the Gulf-Carolinian dataset, Edisto individuals were assigned near-exclusively to a third ancestry component, hereafter ‘Carolinian’ (>95%), EFGB showed the highest representation of the NW Gulf component among Gulf populations, and ancestry proportions exhibited a gradual west-to-east cline characteristic of isolation by distance (**Figure 4H**).

### 3.5 Population co-ancestry

Population-averaged chunk counts (number of RAD loci at which haplotypes share a nearest genealogical neighbor) spanned a narrow range across all site pairs in both datasets (Northern Gulf: 72.2-74; Gulf-Carolinian 65.8-67.6; **Figure 5C, D**), indicating broadly homogeneous shared ancestry among *S. exserta* populations across the WTNWA province. Despite this narrow range, several biologically meaningful patterns were evident. In the Northern Gulf dataset, EFGB showed the lowest mean co-ancestry across all population pairs, with its minimum pairwise value occurring with MTR (72.2; Figure 5C). The highest mutual co-ancestry in the Northern Gulf dataset was observed from McGrail to Elvers (74), followed by McGrail to Bouma (73.9) and RTR to MTR (73.9), indicating proportionally greater shared ancestry among these geographically proximate central and eastern Gulf sites.

**FIGURE 5.**
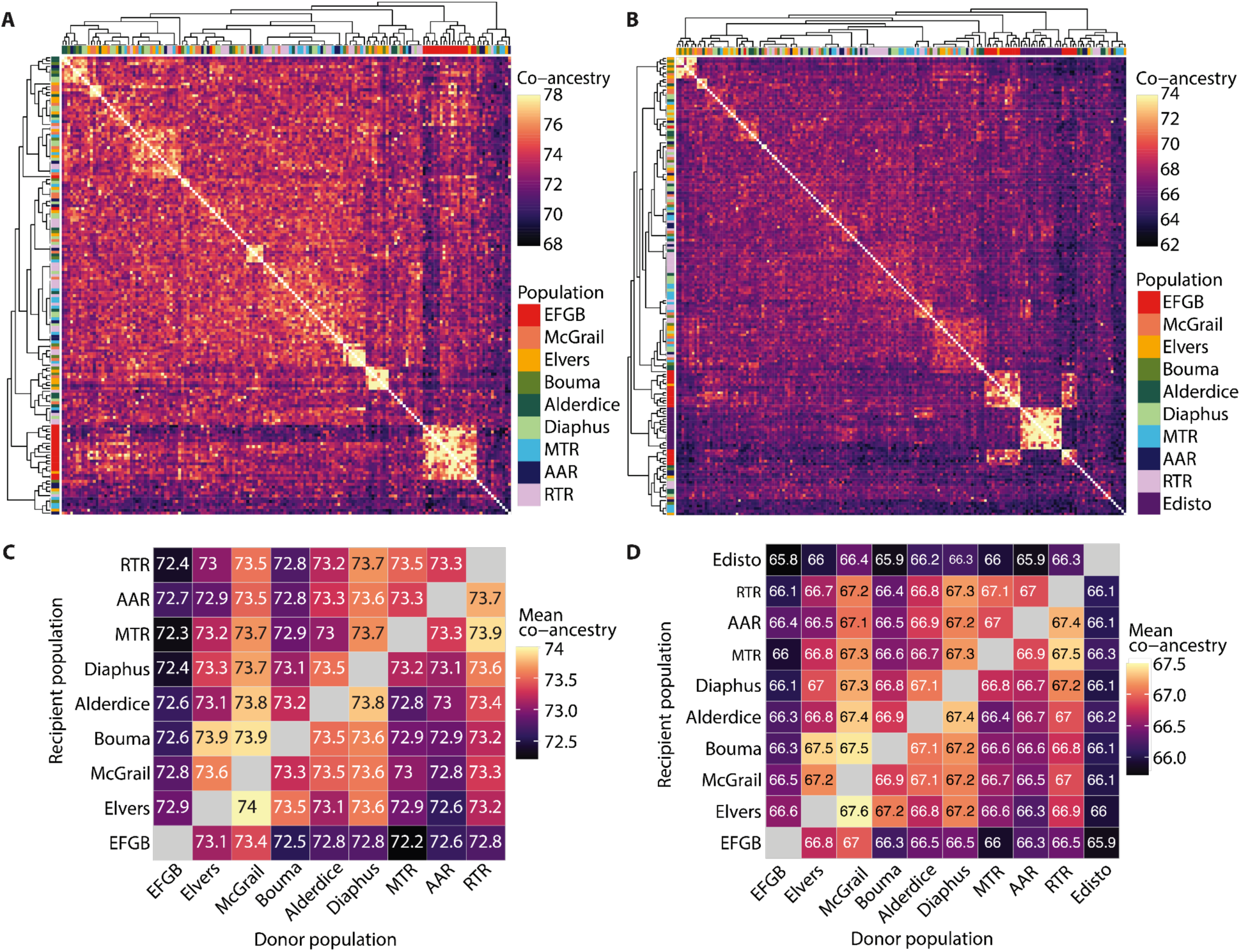
Population co-ancestry and larval dispersal potential of *Swiftia exserta* in the WTNWA province. Individual-level co-ancestry heatmaps of (A) Northern Gulf dataset (9 populations, 158 individuals) and the (B) Gulf-Carolinian dataset (10 populations, 174 individuals). Each cell represents the co-ancestry value between a pair of individuals. Warmer colors indicate greater shared ancestry. Dendrograms along axes reflect hierarchical clustering of individuals by co-ancestry. (C) Population-averaged co-ancestry matrices for the Northern Gulf dataset and (D) Gulf-Carolinian datasets, with rows representing recipient populations and columns representing donor populations.

In the Gulf-Carolinian dataset, Edisto showed lower co-ancestry with all Northern Gulf populations (65.8-66.4, lowest with EFGB) compared to within-Gulf population pairs (66-67.6), confirming its genealogical isolation from the Gulf metapopulation (**Figure 5D**). Within the Gulf, EFGB showed the lowest mean co-ancestry across all Gulf site pairs, with particularly low values relative to eastern populations. Individual-level co-ancestry heatmaps (**Figures 6 A, B**) showed no discrete block structure separating Northern Gulf populations, with off-diagonal values distributed across a narrow and uniform range, supporting continuous gene flow among sites.

**FIGURE 6.**
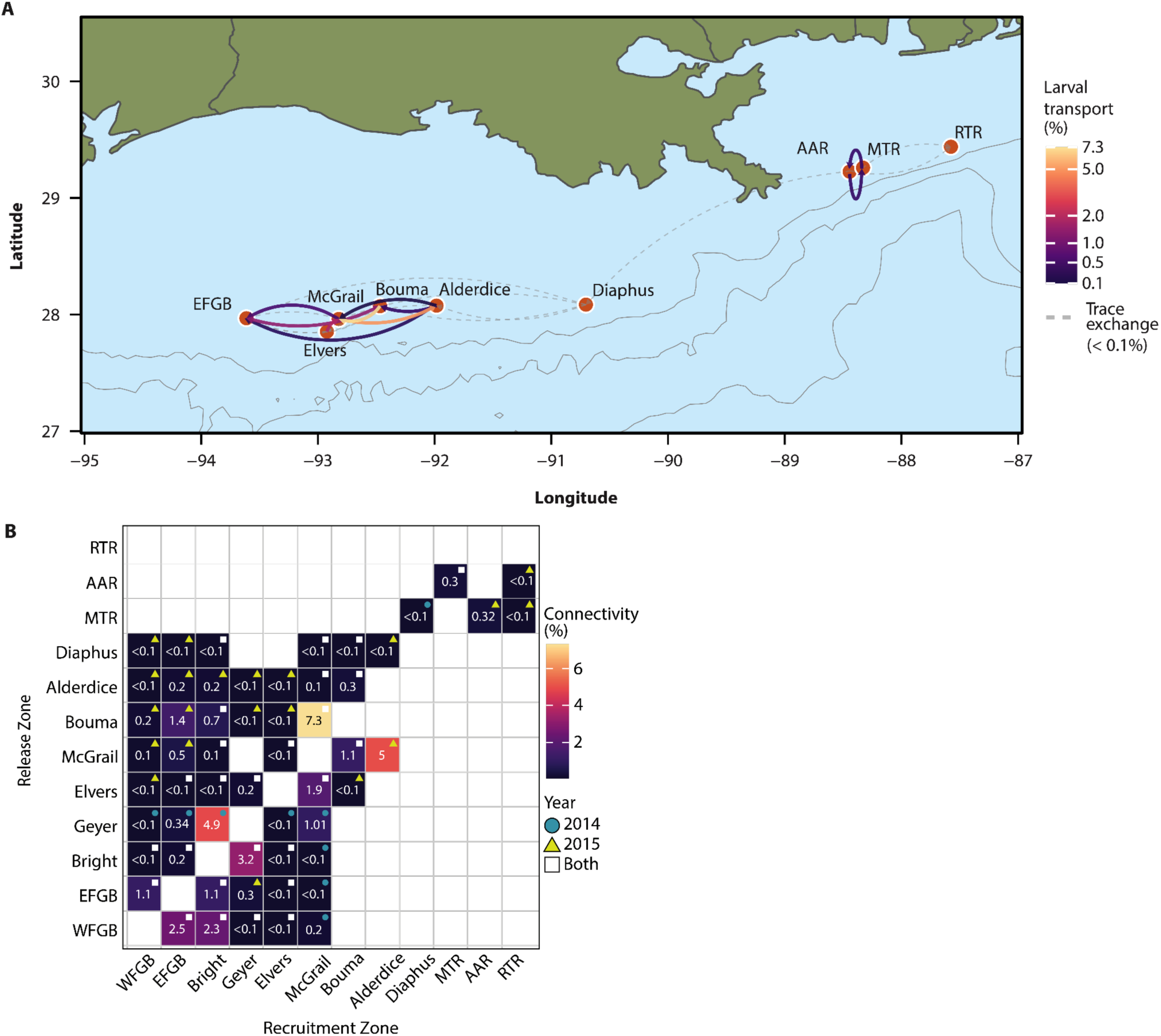
Modeled larval dispersal connectivity among *Swiftia exserta* sampling sites in the WTNWA province, Northern Gulf ecoregion. (A) Arrows indicate the direction of larval transport between sites based on ensemble biophysical particle-tracking simulations during the spawning seasons of September-October 2014 and 2015. Arrow color reflects the percentage of particles successfully transported from a release zone to a recruitment zone across the 60-day simulation period, with darker colors indicating lower transport and brighter yellow indicating higher transport. Dashed grey arrows indicate trace exchange (< 0.1% particle transport) detected in at least one simulation year. Only the nine Gulf populations with ≥ 10 sampled individuals are shown. Contour lines indicate bathymetric depths of 500, 1000, 2000, and 3000 m. (B) Ensemble larval connectivity matrix from biophysical particle tracking simulations during the spawning seasons of September-October 2014 and 2015. Each cell represents the percentage of particles released from a given release zone (rows) that were successfully transported to a given recruitment zone (columns) at any point during the 60-day simulation period. Cell color reflects connectivity strength; symbol shape indicates the year(s) in which connectivity was observed. Populations are arranged west to east along both axes.

### 3.6 Connectivity predicted from physical models

Larval dispersal connectivity was modeled using a hydrodynamical model, CROCO, coupled to a Lagrangian dispersion model, Ichthyop. We simulated two 60-day releases per year at WFGB, EFGB, Bright, Geyer, Elvers, McGrail, Bouma, Alderdice, Diaphus, MTR, AAR, and RTR during September and October 2014 and 2015.

These simulations showed differences in connectivity between 2014 and 2015 (**Figure 6B**). The 2015 simulation predicts a noticeable tendency for eastward connectivity that manifests as a cluster of connections above the diagonal in the connectivity matrix, especially relevant for mid-eastern sites such as Diaphus and Alderdice. Westward dispersal is present, nonetheless, but is not dominant. In contrast, the 2014 simulation, which is representative of ‘normal’ or climatological conditions, exhibits spatially variable connectivity, without a preferential transport direction. Interannual overlap, represented by black cells in the connectivity matrix, is sparse with a limited number of larval exchanges persisting across both 2014 and 2015. This suggests that connectivity in the region is strongly influenced by interannual variations. The strength of the connections is also highly variable, with ensemble values ranging from as high as 7.3% to less than 0.1%. Successful transport of more than 1% of larvae is only predicted in 18% of site connections while fewer than 0.1% is predicted for over half of connections. Overall, the strongest connections are those that occurred during both years, for example between Bouma–McGrail, Bright–Geyer, WFGB–EFGB, and WFGB–Bright.

During the 60-day period considered in our simulations, the sites to the west and east of the Mississippi Canyon appear to be well interconnected within themselves but remain weakly connected between them. Diaphus’s near-solation in the larval dispersal simulations, limited to trace-level exchange (<0.1%) with the NW Gulf cluster and MTR, are at odds with its genetic signal. A similar pattern is evident at RTR, which showed no larval export yet shows equally low F_ST_ from its neighbors. This discrepancy could reflect unsampled or unmodeled stepping-stone link between the NW Gulf and NE Gulf clusters near Diaphus. Larval exchange through such pathways would explain the low genetic differentiation despite no modeled connection. Exploration of additional candidate bank sites and genetic sampling in these intermediate zones would help resolve this. Diaphus acts as a potential source for the Flower Garden Banks while receiving larvae only from MTR (**Figure 6**). Among the sources located west of the Mississippi Canyon, no site dominates, as each of them exported particles to at least five other sites. In contrast, among the sinks, the most recurrent receiving areas span from WFGB to McGrail. In comparison, Alderdice and Diaphus receive from fewer sources and appear as less frequent sinks. Nearly all recruitment zones, from WFGB to Bouma, received particles during both 2014 and 2015, highlighting the steadiness of their role as sinks. The sites located to the east of the Mississippi Canyon appear well interconnected with each other, although RTR did not export any particle during the spawning events considered.

## 4. DISCUSSION

### 4.1 Clonal structure and local population dynamics

Asexual reproduction in *Swiftia exserta* is restricted to individual banks, contributing to local population maintenance but playing no detectable role in inter-site connectivity. All validated genets were confined to single sampling sites, with mean within-group distances of 8 meters, a range previously reported for shallow-water octocorals (Coffroth & Lasker, 1998). While this is the first description of clonality in *S. exserta*, high clonality has been observed in mesophotic scleractinian corals within the region (Drury et al., 2020; Studivan & Voss, 2018b). The tight spatial clustering of genets, and the absence of any cross-site clonal genotype, indicates that asexual propagation in this species operates at sub-decameter scales and does not generate dispersive propagules capable of inter-bank transport. The proportion of sampled individuals belonging to genets should be interpreted cautiously, as non-random ROV-based sampling targeted accessible aggregations rather than employing a systematic spatial design, and clonal prevalence estimates in sessile marine invertebrates are sensitive to sampling strategy (Baums, 2008). Targeted collection of aggregated colonies likely inflates apparent clonality relative to a spatially randomized survey. Nonetheless, the presence of clonal genotypes at 10 of 13 Northern Gulf sites, confirms that asexual propagation is a pervasive feature in *S. exserta* population biology across the banks.

### 4.2 Population genetic structure

*Swiftia exserta* comprises at least two geographically structured lineages, in the Gulf and along the southeastern U.S. Atlantic Coast, with limited gene flow between them; whether these lineages represent independently evolving species remains an open question. Within the Gulf, populations form a single continuously structured metapopulation in which geographic distance drives genetic differentiation. A significant positive relationship between linearized F_ST_ and the natural log of geographic distance provides formal support for isolation by distance as the primary structuring mechanism, validated by positive F_ST_ values across all site pairs, and a continuous west-to-east gradient in ancestry proportions. The absence of hard genetic break across the Mississippi Canyon, despite near-total physical isolation between areas on either side identified by similar model with expanded release scenarios <u>(Lopera et al., 2025)</u>, suggests that the canyon functions as a gradually accumulating soft barrier: even rare larval exchange events are sufficient to prevent genetic differentiation from accumulating into a discrete discontinuity. For deeper octocorals,(e.,g., *Paramuricea biscaya*, *Callogorgia delta)* depth was found to primarily structure connectivity (Galaska et al., 2021). The contrast with *S. exserta*’s distance-mediated pattern may reflect its narrower bathymetric range across the banks.

Within this continuum, EFGB is the most differentiated Gulf population, with the highest pairwise F_ST_ values against all other Gulf sites, the lowest mean co-ancestry in both datasets, and separation along the first DAPC discriminant axis. This pattern reflects EFGB’s position at the western edge of the sampled sites, where it receives proportionally less larval input from the broader Gulf source pool. EFGB’s elevated differentiation, therefore, represents the western extreme of the IBD gradient. While this pattern is consistent with distance-limited gene flow, we cannot rule out a contribution from local environmental differentiation at EFGB, which was not directly tested here, however its depth range falls within that of other Gulf sites. Including the low-sample-size sites excluded from structural analyses, WFGB clusters with EFGB in ADMIXTURE and DAPC, while Bright and Geyer instead pattern with the central Gulf cluster.

The Carolinian ecoregion site, Edisto, is categorically distinct from this Gulf-wide pattern. F_ST_ values between Edisto and all northern Gulf populations substantially exceed the within-Gulf maximum, hierarchical AMOVA identifies the Gulf-Carolinian partition as the largest single source of molecular variance (though not itself statistically significant, likely due to limited power from the single-population Atlantic samples), and Edisto individuals form an isolated cluster in DAPC ordination and individual fineRADstructure heatmaps. *S. exserta* thus adds an octocoral to the set of western Atlantic taxa showing Gulf-Atlantic genetic partitioning across the Florida Peninsula. Comparable Gulf-Atlantic partitioning has been demonstrated in the deep reef-building scleractinian coral *Desmophyllum pertusum* (formerly known as *Lophelia pertusa*). (Morrison et al., 2011) identified regional genetic structure across the Gulf, southeastern U.S. (SEUS), New England Seamounts, and eastern North Atlantic population, although differentiation between Gulf and SEUS populations was relatively weak and included evidence of admixture near southeastern Florida. (Weinnig et al., 2024) found high connectivity across SEUS populations along the U.S. eastern continental margin, while these populations were differentiated from both Gulf populations and populations farther north, including Norfolk Canyon. Interestingly, Norfolk Canyon populations exhibited relatively low genetic differentiation from the Gulf despite their geographic separation, suggesting that connectivity in this species is not explained by geographic distance alone and may reflect the region’s complex oceanographic setting. Similarly, northwestern Gulf populations of the giant barrel sponge, *Xestospongia muta*, differed from all Florida Reef Tract sites (Bernard et al., 2018). Our results extend this pattern into mesophotic octocorals, a group for which taxonomic and regional coverage remains uneven (Banha et al., 2025; Eyal et al., 2021). Sampling populations along the Florida Shelf would allow formal testing of whether Gulf-Carolinian divergence reflects a discrete phylogeographic barrier or the terminus of a steep IBD gradient extended across the Florida Peninsula.

### 4.3 Larval connectivity and its genomic signature

Biophysical larval dispersal modeling and genomic co-ancestry analysis independently recovered the same spatial connectivity structure. The site pairs with the strongest and most temporally stable modeled larval exchange, observed in both 2014 and 2015, McGrail-Bouma and Elvers-McGrail, also exhibited the highest mutual co-ancestry in the genomic data. EFGB’s reduced larval input from the broader Northern Gulf source pool is reflected in its consistently low-co-ancestry values across both datasets. Conversely, EFGB shows the lowest mean co-ancestry across all Gulf site pairs, mirroring the model’s prediction that EFGB receives larvae primarily from nearby western sites, as opposed to the broader Northern Gulf source pool. The forward-modeling larval transport from physical oceanography and historical gene flow inferred from genome-wide SNPs, arrived at the same spatial patterns, strengthening the inferred structure. Similar convergence between particle-tracking models and genomic data has been documented in corals from the Gulf, including studies of *Montastraea cavernosa* and *Paramuricea biscaya* (Garavelli et al., 2018; Liu et al., 2021; Studivan & Voss, 2018a). Comparable integration of genomic and biophysical approaches has also revealed spatially structured connectivity networks in *Acropora milipora* and *Paramuricea biscaya* (Galaska et al., 2021; Matz et al., 2018).

The episodic and temporally variable nature of larval exchange among Northern Gulf banks likely underlies the genomic IBD signature, as variable dispersal accumulated across generations and produced a distance-decay pattern in genetic similarity. Because connectivity depends on the absolute number of successful dispersers per generation, relatively low rates of gene flow can reduce genetic differentiation among otherwise demographically independent populations, even when those migrants make only a small contribution to local recruitment (Lowe & Allendorf, 2010). IBD-based dispersal estimates have been validated directly against genetic parentage assignments in a Caribbean reef fish, recovering nearly identical dispersal kernels (Naaykens & D’Aloia, 2022). The genetic signal integrates dispersal and gene-flow processes operating over multiple generations, whereas biophysical models characterize connectivity under specific oceanographic conditions and time periods (Raynal et al., 2014; Selkoe & Toonen, 2011).

Thus, any single year of biophysical modeling represents only one realization of the broader distribution of connectivity outcomes, and the agreement observed here reflects the correspondence between genetic connectivity accumulated over evolutionary timescales and the longer-term tendency of physical connectivity, with contemporary larval flux representing a shorter-term realization of that process (Chérubin & Garavelli, 2016; Holstein et al., 2014; Kough & Paris, 2015). Genomic and biophysical approaches are therefore complementary, and multi-year or ensemble modeling is preferable to single-season estimates for inferring ecologically relevant connectivity (Garavelli et al., 2018; Naaykens & D’Aloia, 2022).

### 4.4 Broader implications

The IBD-structured connectivity corridor from WFGB through McGrail identifies a functionally continuous corridor in the Northern Gulf that should be managed as an interconnected unit. EFGB’s peripheral position and reduced genetic exchange suggest it would benefit disproportionately from management actions that only maintains gene flow from the central corridor. More broadly, this dual reproductive strategy of site-restricted asexual fragmentation sustaining local density between spawning events, paired with broadcast spawning maintaining genetic cohesion across the region constitutes a regional control on metacommunity dynamics in this system. Site-restricted clonality means colony counts are an inadequate proxy for population genetic diversity, and outplanting from few donor colonies risks establishing low-diversity stands with reduced adaptive potential. As *S. exserta* is a priority species for active restoration, it is imperative that restoration planning ensures genetic diversity when selecting colonies for transplanting. This has been demonstrated in shallow-water coral studies, where accounting for genotypic diversity among outplants improves performance metrics including growth and reproductive output (Baums et al., 2019; Rios et al., 2025). The slow growth rates, habitat patchiness, and disturbance exposure that characterize mesophotic coral ecosystems in the Northern Gulf make the interplay between local clonal resilience and regional sexual connectivity particularly consequential for long-term metacommunity persistence.

## AUTHOR CONTRIBUTIONS

S.H., A.M.Q., and A.B. conceptualized the study and acquired funding. S.H. M.P.G. and D.W. collected samples and generated sequencing data; N.C.P., S.H., L.L. and S.A.V. analyzed the data; N.C.P., S.H., and L.L. wrote the first version of the manuscript; all authors revised the manuscript and approved the final version.

## ACKNOWLEDGMENTS

We thank the captains and crew of expeditions OP17 (MSV Ocean Project), MT17 and MT18 (R/V Manta), PE19-25 (R/V *Pelican*) and PS21-04 (R/V *Point-Sur*), as well as ROV pilots and technicians from Oceaneering Inc. and UNCW’s Undersea Vehicles Program. We are particularly thankful for the contributions of Jill McDermott, Alexis Weinnig, Janessy Frometa, Fanny Girard, Amanda Glazier, Alondra Maldonado, Juan Sanchez, Luisa Dueñas, Peter Etnoyer, Maria Granquist, Cathy McFadden, Cheryl Morrison, and Frank Parker to this project. Special thanks to the Flower Garden Banks National Marine Sanctuary staff (G.P. Schmall, Emma Hickerson, Marissa Nuttall, and Michelle Johnston) for their support and engagement.

## PERMITS

Work within the Flower Garden Banks National Marine Sanctuary was conducted under permits FGBNMS-2017-007-A2 and FGBNMS-2019-003-A2 to S.H.

## DATA AVAILABILITY STATEMENT

*Swiftia exserta* raw sequence reads are available at BioProject PRJNA821015 (https://www.ncbi.nlm.nih.gov/bioproject/PRJNA821015). The whole genome assembly has been deposited at DDBJ/ENA/GenBank under the accession JCANKD000000000. The version described in this paper is version JCANKD010000000. Associated datasets and data tables are available at https://doi.org/10.6084/m9.figshare.33292446 and code at https://github.com/npittoors/Swiftia_exserta_GoM_PopGen.

## SUPPORTING INFORMATION

**Supplementary Figure 1.**
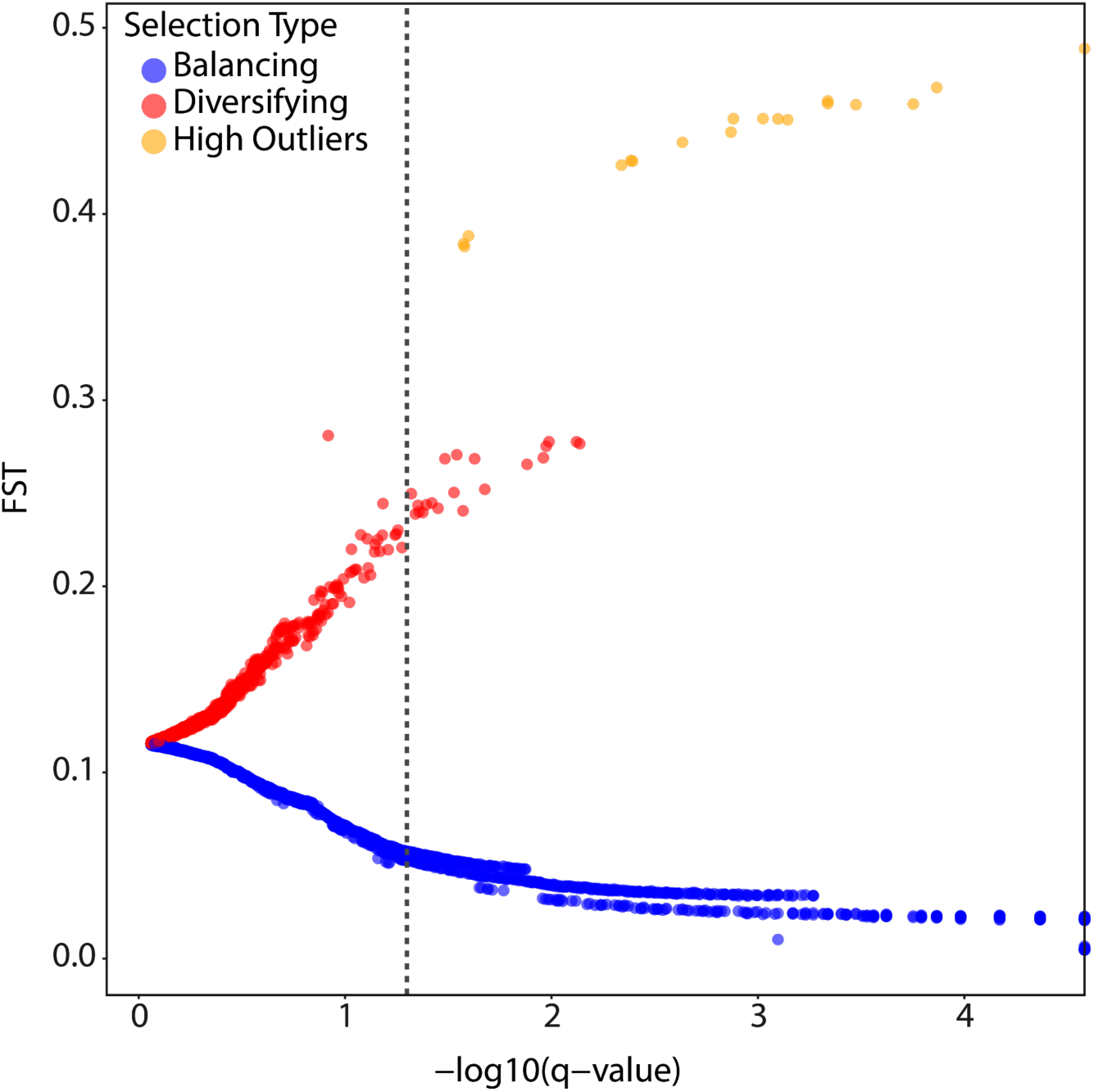
BayeScan outlier detection results for *Swiftia exserta* SNP loci. Each point represents a single SNP locus plotted by posterior odds of selection (-log10(q-value), x-axis) against locus-specific F_ST_ (y-axis). Points are colored by selection classification balancing selection (blue, negative alpha values), diversifying selection (red, positive alpha values, log10(PO) > 2), and high-confidence outlier loci retained for removal (orange, F_ST_ > 0.35, q < 0.05, n = 18). The dashed line indicated the significance threshold (-log10(q-value) = 1.3, corresponding to q = 0.05).

**Supplementary Figure 2.**
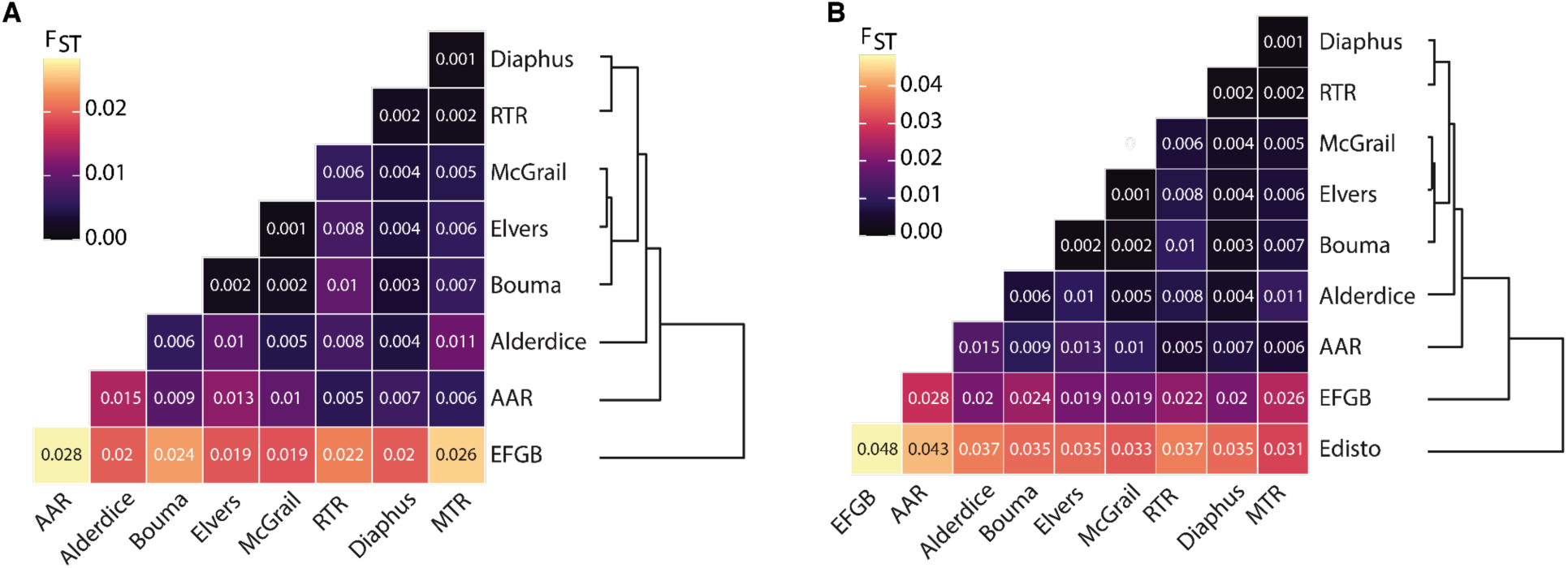
Pairwise F_ST_ among *Swiftia exserta* populations in the WTNWA province, ordered by hierarchical clustering. F_ST_ values for (A) the Northern Gulf dataset and (B) Gulf-Carolinian dataset. Populations are arranged to hierarchical clustering of the F_ST_ distance matrix, with the corresponding dendrogram shown to the right.

**Supplementary Figure 3.**
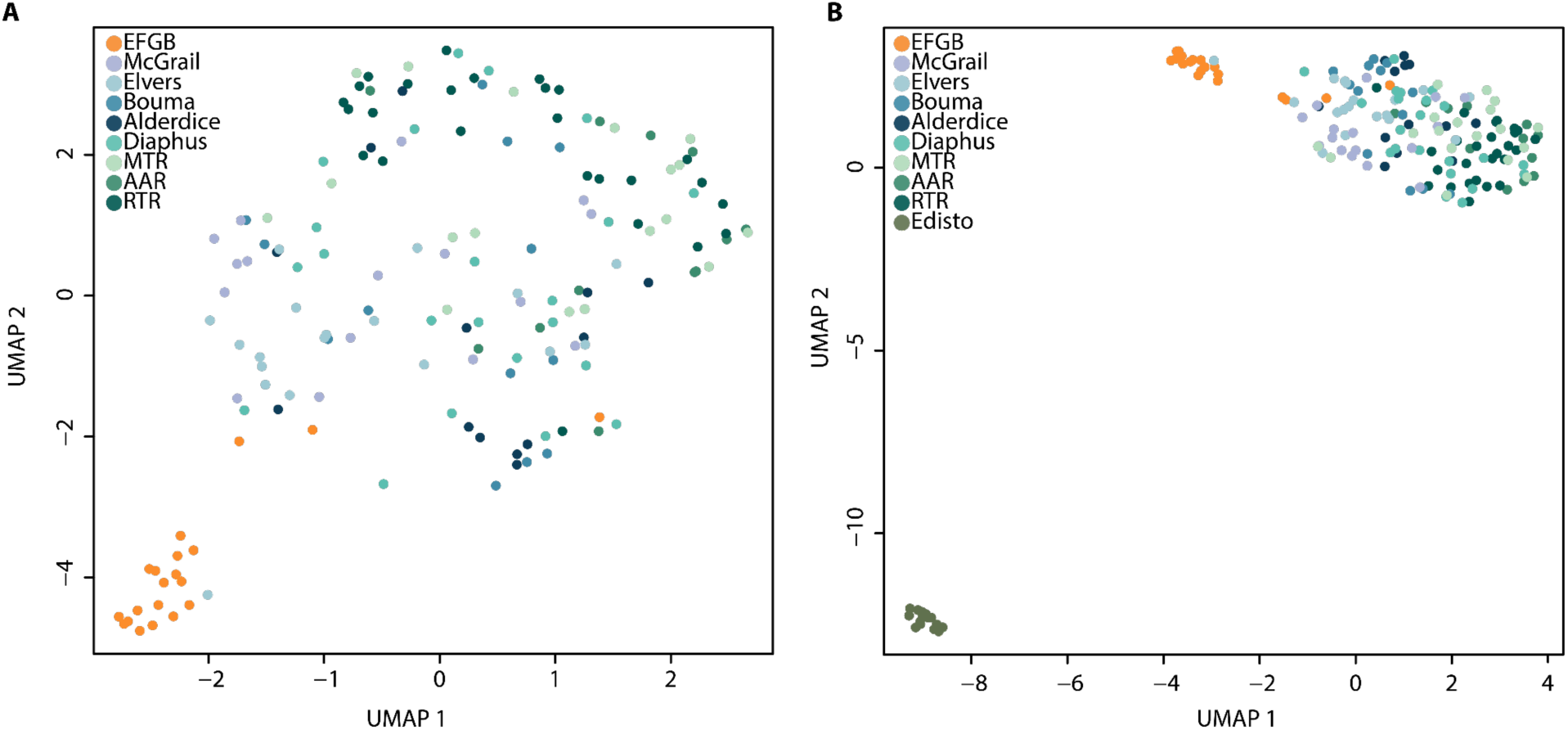
UMAP ordination of DAPC discriminant scores for *Swiftia exserta* populations in the WTNWA province. (**A**) Northern Gulf dataset and (**B**) Gulf-Carolinian dataset.

